# Dynamics of oscillatory power subtends multiplexing of perceptual and mnemonic information within the premotor cortex

**DOI:** 10.1101/2025.05.08.652817

**Authors:** Isabel Beatrice Marc, Valentina Giuffrida, Surabhi Ramawat, Giampiero Bardella, Stefano Ferraina, Emiliano Brunamonti

## Abstract

Flexible behaviour requires decision-making that integrates both perceptual and mnemonic-related information. While the dorsal premotor cortex (PMd) is known to support decision-making based on perceptual computations, its contributions to decisions based on mnemonic information for motor planning are underexplored. Here, investigating local field potential (LFP) oscillations in PMd of monkeys performing a transitive inference task required the formation and retrieval of mnemonic representations of an arbitrarily defined rank order among perceptual items. Our results highlight that a dynamic interplay between lower frequencies (Theta, Alpha, Beta) and high-Gamma oscillatory activity of LFP reflects a mechanism for accessing and manipulating memory-related information underlying decision-making. These findings provide evidence that the PMd plays a role in multiplexing both perceptual and mnemonic information, extending its competence beyond the association of perceptual input with motor decisions.

## INTRODUCTION

In our interaction with the environment, we continuously decide among alternatives. This form of decision process has been routinely studied at the neural level in animals solving a task using explicit information, sometimes cognitively demanding, mostly proposed through the visual system^1–4^. The PMd, a key region involved in sensorimotor transformations and the use of learned arbitrary stimulus-response associations^5,6^, plays a crucial role in different forms of decision-making processes that lead to motor actions^3,7,8^. It is known to encode motor plan maturation and multiple motor options^7^, even when the movement is not subsequently executed^9^. Furthermore, neurons in PMd adjust their firing rate activity based on decision difficulty, whether driven by perceptual^2,3^ or memory-schema ambiguity^10,11^.

Thus, PMd emerges as a key component at the interface between perceptual and abstract information processing for action generation^12^. Here, we investigate the underlying mechanisms by examining neural oscillation modulations through a transitive inference (TI) task^13^. The different forms of TI task previously used in animals and humans demonstrate the brain’s ability to implicitly assign ordinality to pictorial items based on independently learned relationships^13–15^. By experiencing pairwise associations (e.g., A>B, B>C, C>D, etc.), subjects construct a hierarchical mental representation (e.g., A>B>C>D>E>F), enabling them deducing transitively relationships between items never directly compared (e.g., B>D). To determine the higher-ranking item in a pair, subjects must access, during each trial, the individual stored representation, retrieve and temporarily reactivate the ranking information specific to the items under comparison, engaging at the same time both perceptual and mnemonic processes. This suggests that the neuronal networks supporting this form of decision should multiplex explicit perceptual and implicit mnemonic information to drive decision deliberation. Here, we show that TI computation could be described in how LFP oscillations are reciprocally modulated in PMd, providing insight into how incoming information is processed within the network^16^. As monkeys solve the TI task, PMd displays a significant modulation in opposite directions of low (Theta, Alpha, Beta) and high (Gamma) frequency bands, depending on the learned relative rank of items to be compared. Fitting within a model where the transition from information processing to decision deliberation is reflected in similar processes in other areas^16,17^, we propose that the reported LFP modulations reflect how incoming perceptual and mnemonic information are dynamically multiplexed in PMd.

## RESULTS

### Evidence of manipulation of a mnemonic linear representation of ranked items

We tested two monkeys (Macaca mulatta) on a TI task in which a set of six items were arbitrarily rank-ordered (color-coded and labelled as ‘A’, ‘B,’ ‘C,’ etc.; in Fig. 1**a**). The monkeys were required to compare pairs of items presented simultaneously on a computer monitor after learning their relative ranks through a learning session that involved comparing pairs with adjacent ranks (Methods and Fig. 1**a-d**). The symbols used and their position were continuously changed across sessions. To determine the higher-ranking item in a pair, the monkeys must perceptually decode each item and match the perceptual information to the stored mnemonic representation of its relative rank acquired during learning (Fig. 1**d-e**). Deciding within this context could vary in difficulty in function of the difference in rank between the items, emerging as a Symbolic Distance (SD) effect (Fig. 1**c**). This is because the mnemonic representation of rank order is hypothesized to be linearly organized, with items of similar rank positioned adjacently in the representation^13,15,18^.

**Fig. 1.**
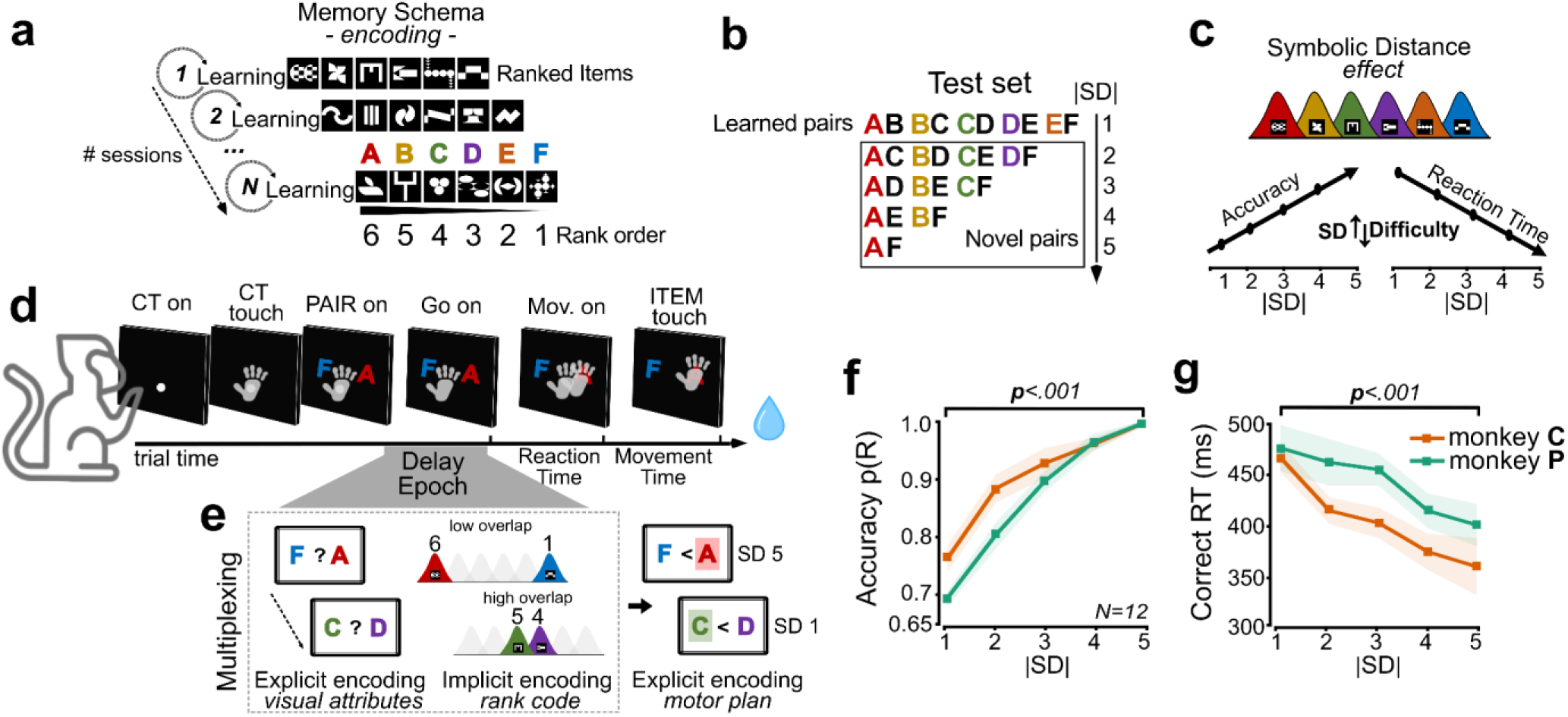
Transitive inference (TI) task and behavioural performance b. *Experimental design*. **(a-d). a** Rank hierarchy of six items (A–F), color-coded for clarity. Training involved adjacent pairs (one-rank difference, SD=1), with rewards given for selecting the higher-ranked item. The ranked series has been changed for each recording session. **b** Testing phase included both learned (adjacent) and novel (non-adjacent) item pairs (e.g., D < B) to assess the monkey’s ability to rank order generalization. The six-item set yielded 5 SDs and 15 pair comparisons, with higher-ranked item positions randomly counterbalanced between left and right. **c** Hypothesized spatial representation of the items’ rank hierarchy. The SD effect is illustrated through item distributions, where lower SDs cause greater overlap in mental representation, making comparisons harder, while higher SDs reduce overlap, facilitating decisions. **d** Trial structure: Monkeys touched and held a central target (CT) before a pair of items appeared (PAIR on). After a variable delay, a Go signal (Go on) prompted them to release the target (Mov. on) and select one of the two items (ITEM touch). **e** During the delay epoch, we hypothesize a multiplexing process that develops in 2 sequential stages: first, an explicit encoding of the visual attributes of the items in the pair; second, through this visual encoding, an implicit encoding of each item’s rank code information. After the Go signal is presented, this information becomes the basis for an explicit encoding of the decision into a motor plan. **f-g** Accuracy p(R) and Correct RTs as functions of SDs across sessions (*N*=12; performed by each monkey). Shaded areas indicate ±1 SEM.

According to this framework, we detected that as SD increases, the decision Accuracy improves and the Reaction times for correct trials (Correct RTs) decline in both monkeys (Fig. 1**f-g**; Supplementary Fig. 1a-b and Table **S**1 for detailed session information; one-way repeated measure ANOVAs). Specifically, for monkey C (Accuracy: *F* _(4,44)_ = 32.09, p < .001; RTs: *F* _(4,44)_ = 11.99, p < .001; sessions *N*=12) and monkey P (Accuracy: *F* _(4,44)_ = 106.35, p < .001; RTs: *F* _(4,44)_ = 10.16, p < .001; sessions *N*=12). The spatial location of the higher-ranked item was not significantly different (see Supplementary Fig. 1c-e; two-way repeated measures ANOVAs, p > .05). With this evidence in hand, we proceeded to investigate how neural activity in the PMd could explain the hypothesized mechanisms underlying this form of decision-making.

### Rank code comparison difficulty modulates the power of lower frequency and high-Gamma bands

To assess how rank code comparison difficulty is reflected in LFPs, we analysed oscillatory power across four frequency bands: Theta (4–7.5Hz), Alpha (8–13Hz), Beta (14–30Hz), and high-Gamma (65-120Hz) bands in the epoch starting when items are presented (PAIR onset) and animals are computing their relationship (delay epoch; Fig. 1**d-e**), multiplexing perceptual and mnemonic information. LFPs were recorded from a Utah array implanted in the left PMd of both monkeys, contralateral to the performing arm (see Methods; Fig. 6**a-b**). LFPs capture synaptic and dendritic activity underlying input integration and are commonly linked to cognitive network computations, offering insight into how neural circuits coordinate during complex tasks^19,20^. While lower frequency oscillations regulate input processing, high-Gamma activity, in contrast, largely reflects spiking activity of local cortical ensembles^21–23^. The temporal evolution of oscillatory power following the PAIR onset revealed a change in power as a function of the difficulty of the decision-related processes in distinct phases. In both monkeys, lower frequency bands showed an initial increase (∼200ms) in power immediately after PAIR onset. However, this initial increase was followed by a power decrease modulated by rank code comparison difficulty (Fig. 2**a-c**). Contextually, this modulation was mirrored by the increasing power in the high-Gamma band (Fig. 2**d**). The observed modulations of the decision-related activity were statistically significant both for the lower frequencies and high-Gamma bands (one-way ANOVAs, p < .001; Bonferroni corrected with **α**=.01). Small insert panels illustrate the averaged channel-modulated periods across different SDs, showing that averaged lower frequency power modulation gradually decreased with increasing SD, while high-Gamma power increased with higher SD (one-way repeated measures ANOVAs, p < .001; see Supplementary Table S2).

**Fig. 2.**
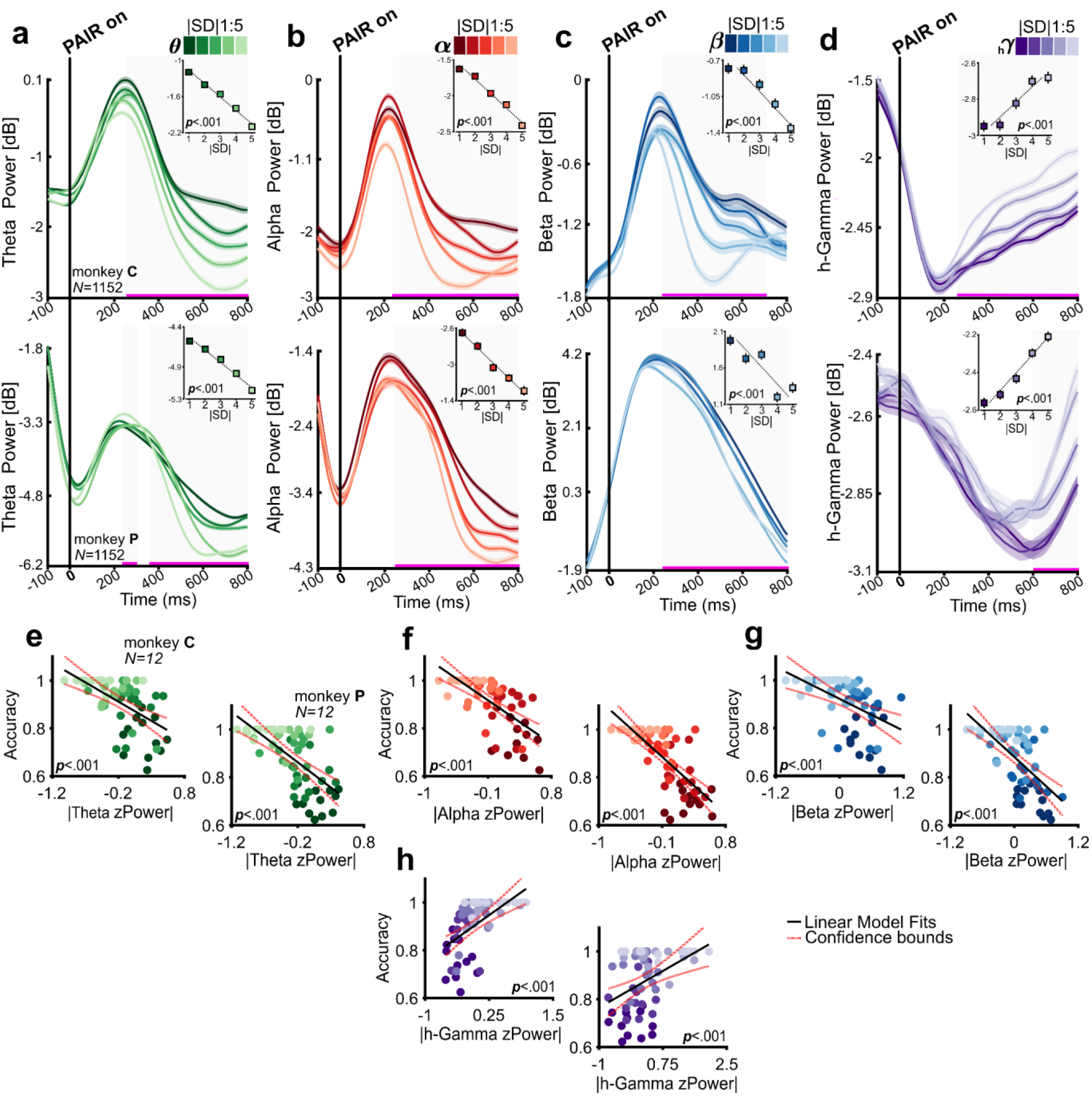
SD-modulations of lower frequency and high-Gamma bands power aligned to PAIR onset. **a-d** Theta power (4-7.5 Hz; **θ** - shades of green), Alpha power (8-13 Hz; **α** - shades of red), Beta power (14-30 Hz; **β** - shades of blue), and high-Gamma power (65-120 Hz; **hγ** - shades of purple) across different SDs (1 to 5; dark to bright colours). The upper panels show results for monkey C and the lower panels for monkey P, averaged over channels, *N*=1152 each, with shaded areas indicating ± 1 SEM. Significant differences in time are marked with a solid pink line and a grey background in each plot (one-way ANOVAs, p < .001, Bonferroni corrected with **α** = 0.01). The x-axis spans from −100 ms before to +800 ms after the PAIR onset. The y-axis indicates power magnitude in decibels (dB), normalized to the baseline period. Averaged power across all channels for each SD, shown separately (inserts) for each monkey (one-way repeated measures ANOVAs, all ps < .001). **e-h** Scatter plots showing the relationship between normalized power (zPower) and Accuracy across sessions and SD (12 (sessions) x 5 (SDs) for each monkey). Linear regression analyses revealed a negative relationship for lower frequency bands and a positive relationship for high-Gamma (p < .001). The left panels show results for monkey C, and the right panels show results for monkey P. The linear regression model fits (black solid lines) and confidence bounds (dotted red lines) are illustrated.

Furthermore, linear regression analyses revealed that the observed modulation in lower frequencies was negatively related to Accuracy (monkey C: Theta slope −0.16, p < .001; Alpha slope −0.20, p < .001; Beta slope **-**0.11, p < .001; and monkey P: Theta slope **-**0.15, p < .001; Alpha slope **-**0.16, p < .001; Beta slope **-**0.19, p < .001) (Fig. 2**e-g**). In contrast, the high-Gamma band modulation was positively related to Accuracy (Fig. 2**h**) (monkey C: slope +0.15, p < .001; monkey P: slope +0.10, p < .001). Overall, these results align with the complementary role of the lower frequencies and high-Gamma bands in decision-making, where the change in power of high-Gamma is associated with decision deliberation occurring alongside opposite changes in the power of lower frequencies^16^. The gradual increase in high-Gamma power as difficulty decreases, accompanied by a corresponding decrease in lower frequencies’ power, suggests that the decision-making process is supported by this dynamic interplay between frequency bands.

### The high-Gamma band differs in magnitude interplaying with different lower frequency bands

Building on our earlier findings of SD-related changes scaling with rank code comparison difficulty, we further examined the relationship between all lower frequency bands and high-Gamma. Two key features emerged: **i**) an inverse relationship between the power of lower frequencies and high-Gamma, and **ii**) a modulation of this inverse trend by rank code comparison difficulty (see Fig. 2**a-h**). To quantify the strength of this inverse modulation of each lower frequency with high-Gamma, we performed cross-frequency Pearson correlation analyses at the channel level (Fig. 3**a**). A negative coefficient at a given time point indicates that, across SDs, reduced power in a lower frequency band was associated with higher power in the high-Gamma band. We found that this anticorrelation significantly emerged across the great majority of channels and for both monkeys (Fig. 3**b**). A comparison of anticorrelation peaks strength in both monkeys (Fig. 3**c**) revealed significant differences in frequency band interactions (one-way repeated measures ANOVAs using channels *N*=96: monkey C, F _(2,190)_ = 18,16, p < .001; monkey P, F _(2,190)_ = 23,56, p < .001).

**Fig. 3.**
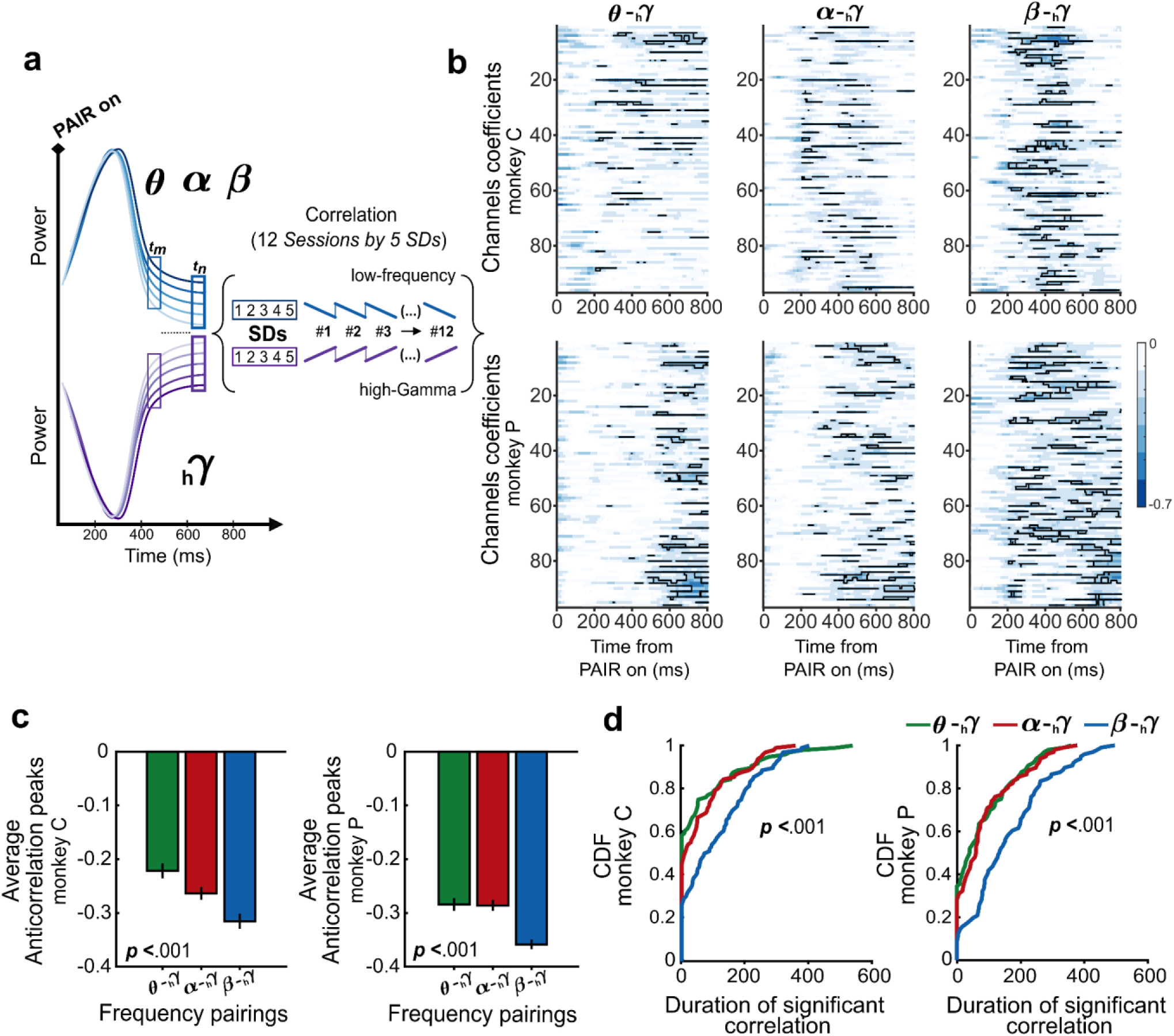
Temporal interplay of lower frequencies and high-Gamma band power across channels. **a** Schematic of cross-frequency correlation analysis. For each channel, power values in lower frequencies (θ, α, β) and high-Gamma (hγ) were extracted at each time point (*tn*) during the delay epoch. Power values from all sessions and SDs were collected separately for each frequency band, resulting in two time series per channel: one for each lower frequency (blue) and one for high-Gamma (purple). Pearson correlation coefficients were calculated between these time series at each time point, generating a cross-frequency correlation series for each channel and frequency pairing. **b** Cross-frequency correlation time series for all channels in the epoch of reference (θ–hγ, α–hγ, β-hγ; left to right panels). Based on prior identification of modulation periods (from 200ms after PAIR onset to 800ms) statistically significant correlations within this window are indicated by black lines contours. **c** Average anticorrelations peaks across all channels for each frequency pairing, displayed as bar plots: θ–hγ (green), α–hγ (red), and β–hγ (blue), with error bars representing ±1 SEM. The β–hγ pairing showed the higher anticorrelation overall, significantly differing from the other pairings in both monkeys (one-way repeated measures ANOVAs, p < .001). **d** Quantification of channel-wise significance across time. For each channel the duration of significant time points within the window was computed. Cumulative distribution functions (CDFs) of these counts are plotted, illustrating the distribution of temporal significance across channels and highlighting the prevalence of sustained cross-frequency anticorrelation during the delay epoch among the β–hγ pairing (one-way repeated measures ANOVAs, p < .001).

Post hoc tests confirmed that the **β** – **hγ** pair showed a significantly stronger anticorrelation compared to both **θ** – **γh** (monkey C: p < .001; monkey P: p < .001) and **α** – **hγ** (monkey C: p = .002; monkey P: p < .001), while the difference between **θ** – **hγ** and **α** – **hγ** was significant only for monkey C (p =.01) but not for monkey P (p > .05). Moreover, a quantification of duration of the significant anticorrelations (Fig. 3**d**) revealed that it was longer in both monkeys for **β** – **hγ** than for **θ** – **γh** or **α** – **hγ** (one-way repeated measures ANOVAs monkey C, F _(2,190)_ = 8.77, p < .001; monkey P, F _(2,190)_ = 23,29, p < .001). The varying degree of correlation between the power of different lower frequencies with the high-Gamma suggests a distinct contribution of the lower frequency bands to the decision deliberation process. The stronger correlation observed between the decrease in Beta power and the increase in high-Gamma power indicates a closer functional relationship between these two frequency bands compared to the relationship between high-Gamma and either Theta or Alpha. Moreover, this analysis, conducted at the channel level across multiple sessions with different rank-ordered items, reflects consistent neural computations associated with rank code comparison difficulty rather than stimulus perceptual features, which vary across sessions.

### Channel recruitment is influenced by the rank code comparison difficulty in PMd

To investigate how rank code comparison difficulty modulates the spatial pattern of lower and high-Gamma bands power across channels, we mapped the SD-related power changes, derived from the significant time periods (Fig. 2**a-d**) and averaged across sessions, onto the 96-channel Array. Our findings revealed a selective recruitment of channels, consistent with our previous temporal analysis. Specifically, low SDs (difficult comparisons) were associated with greater recruitment of channels and a relatively higher power of lower frequencies, while higher SDs (easier comparisons) showed a relatively low recruitment of channels. This modulation was mirrored by a corresponding increase in high-Gamma recruitment of channels and power. These spatial modulations suggest that rank code comparison difficulty corresponds to a different involvement of the explored network (Fig. 4**a-b**).

**Fig. 4.**
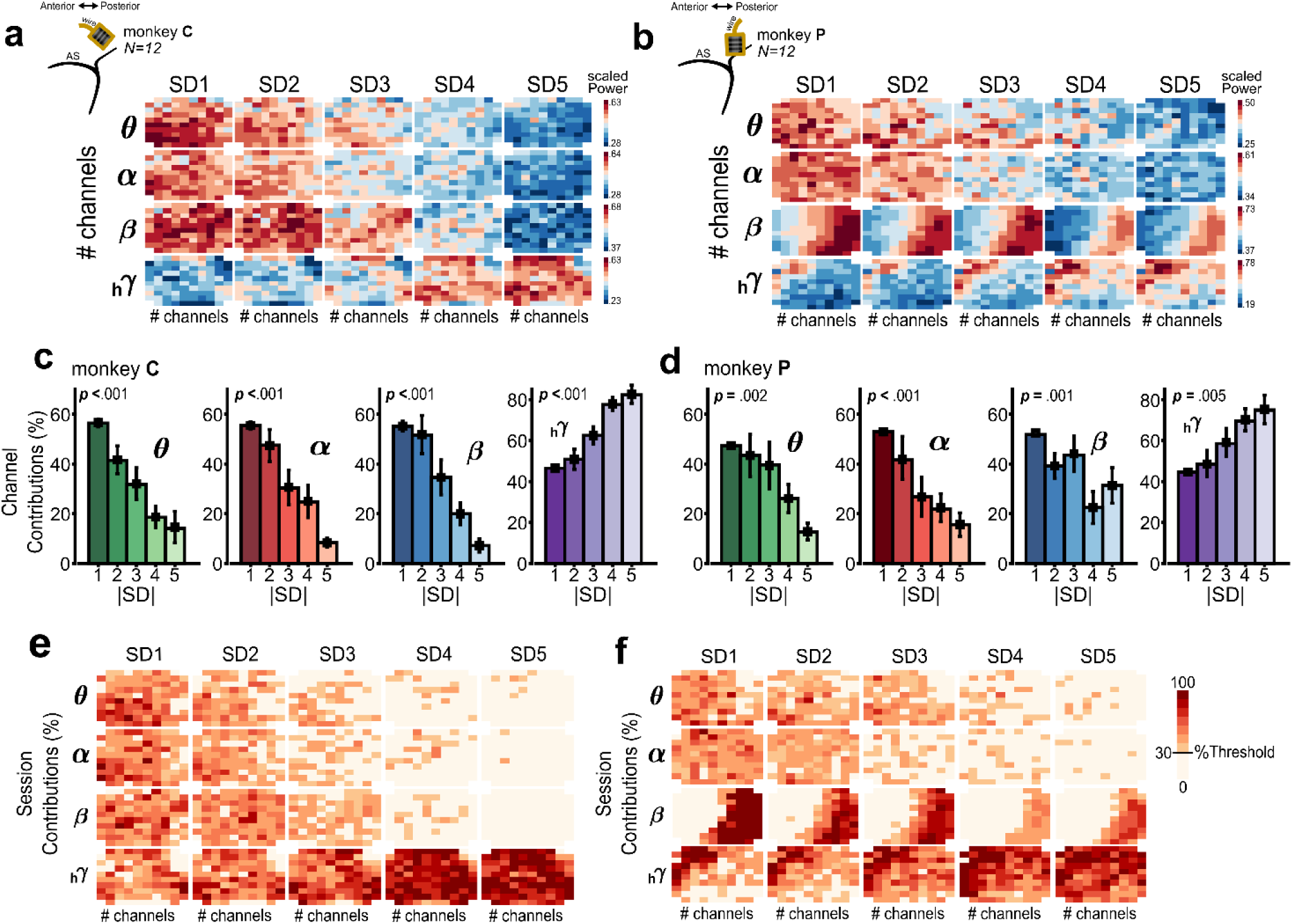
Spatial distribution of channel contributions to LFPs modulation. **a-b** Electrode array placements for monkeys C and P are shown, with the anterior sulcus (AS) and wire positions marked to visualize power distribution across PMd. The channel maps illustrate the spatial extension of SD-modulated power across the 96-channel array, averaged across sessions. Maps are organized by frequency band (θ, α, β, hγ), with each row representing a different frequency band and each column corresponding to SD (from 1 to 5). The colour scale represents scaled power values (0:1), with separate colour bars for each frequency band. **c-d** Bar plots showing the average percentage of channels exceeding the contribution threshold for each frequency band and SD, illustrating how channel contributions (%) vary with rank code comparison difficulty. The plots are color-coded by frequency band (θ, α, β, hγ from green to purple) and show the proportion of significantly contributing channels for each frequency band across sessions for monkeys C and monkey P. Black error bars represent ±1 SEM. **e-f** Channel maps showing the percentage of sessions in which each channel contributed to the SD-modulated power across frequency bands (**θ** to **hγ**) and SD conditions (1 to 5). Each map represents the proportion of sessions in which a given channel exceeded the contribution threshold, with colour coding reflecting the percentage of sessions for each channel. A colour bar indicates the percentage of sessions contributing across channels, with scaling based on a channel contribution threshold of 30%.

To investigate this further, we calculated each channel’s contribution by comparing its power during significant time periods of the delay epoch within each session to the total power in the reference condition (SD=1). Channels contributing more than 1.04%, a contribution ratio threshold derived from the assumption of equal contribution across all channels (details in the Statistics section), were classified as significant. Results showed that lower frequencies exhibited greater channel percentage recruitment during more difficult decisions (SD=1-2), with a progressive decrease in the number of contributing channels as rank code comparison difficulty decreased (SD=4-5) (Fig. 4**c-d**). Significant effects were observed for both monkeys: for monkey C (Friedman’s tests, Theta: χ² _(4, 44)_ = 27.38, p < .001; Alpha: χ² _(4, 44)_ = 39.27, p< .001; Beta: χ² _(4, 44)_ = 34.52, p < .001) and for monkey P (Friedman’s tests, Theta: χ² _(4, 44)_ = 16.27, p = .002; Alpha: χ² _(4, 44)_ = 23.58, p < .001; Beta: χ² _(4, 44)_ = 17.14, p = .001). In contrast, the high-Gamma showed fewer contributing channels during the most difficult pair comparisons, with this number increasing as rank code comparison difficulty decreased (Friedman’s tests, monkey C: χ² _(4, 44)_ = 36.77, p < .001; monkey P: χ² _(4, 44)_ = 14.83, p = .005). To complete these findings, we quantified whether the channels that were significantly contributing and recruited also showed consistency across sessions (Fig. 4**e-f**). This consistency indicates that the same neural resources were reliably engaged, suggesting stable recruitment of specific channels across sessions depending on the comparison difficulty, which seems to be independent of the pictorial features of the stimuli that continuously change. This complements our previous analysis of the interplay between lower frequencies and high-Gamma power changes, showing that rank-code comparison difficulty modulations remain consistent at the channel level, highlighting the network engagement in processing the cognitive aspects of the task.

### Power characteristics of frequency bands power in Wrong Choices

Having established that the PMd’s LFP oscillations power dynamics in correct choice trials play a role in the processing memory memory-related information subtending decision making, we further investigated how these dynamic changes occur in wrong choice trials. Wrong choices were defined as instances in which the monkey selected the lower-ranked item in a pair. Given the higher response accuracy for simpler comparisons (Fig. 1**f**), wrong trials from SD4 and SD5 (referred to as SD4/5 for simplicity) were combined to increase the statistical power of the following analysis. We calculated RTs for wrong choices in a subset of sessions where Wrong RT were detected in all SDs (monkey C: *N*=7/12; monkey P: *N*=6/12). We then quantified the RT discrepancies by computing a *Difference Index* (Wrong RTs - Correct RTs) for each SD (Fig. 5**a**). Figure 5 shows that Wrong RT was consistently longer than Correct RT for both monkeys (all > 0), with a greater difference in easier comparisons (SD4/5). A two-way ANOVA (Monkey and SD) revealed a significant main effect of SD (F _(1,3)_ = 4.50, p = .007), with post-hoc tests confirming significant differences between SD4/5 and the other (p < .05). No significant main effect of Monkey (*F* _(1,1)_ = 3.29, p= .07) or interaction (*F* _(3,48)_ = 0.28, p = .83) was observed, indicating similar RT patterns across SDs for both monkeys.

**Fig. 5.**
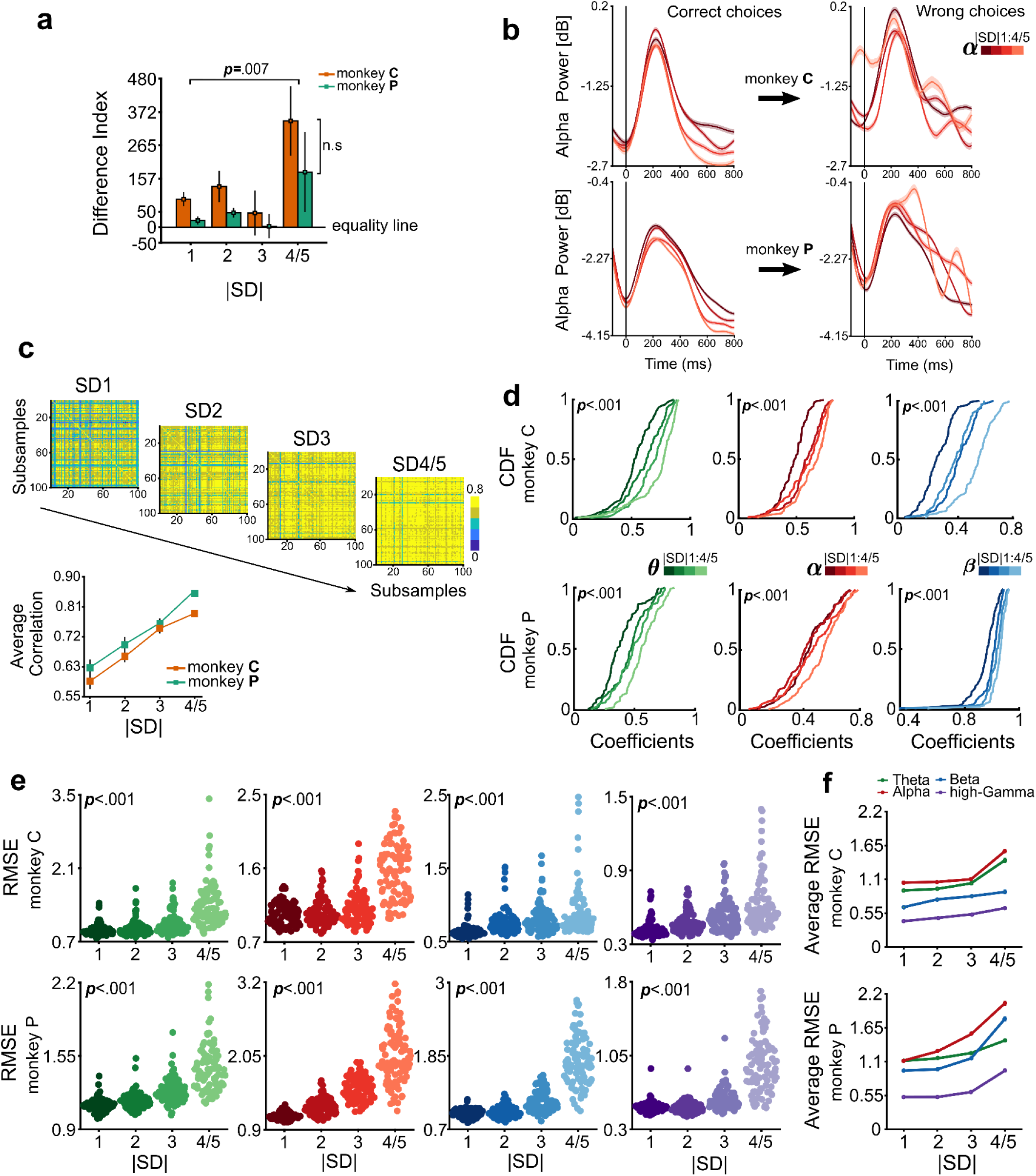
Power dynamics and behaviour in Wrong choice trials. **a** Difference Index (Wrong RT - Correct RT) for monkeys C (orange) and monkey P (green) across available sessions (monkey C: *N*=7/12; monkey P: *N*=6/12). Error bars indicate mean ± 1 SEM. **b** Example of Alpha power (shades of red) aligned to PAIR onset for monkey C (upper panels) and monkey P (lower panels) across SDs (1 to 4/5). Power dynamics are compared between correct and wrong trials, showing increased variability during wrong choices. **c** Correlation matrices for a single channel across 100 randomly selected subsamples of correct trials for each SD condition, showing the similarity of power dynamics during modulated time periods. The lower panel shows the average correlation for one channel per monkey, with error bars indicating ± 1 SEM. **d** Channels Cumulative distribution function (CDF) of Pearson correlation coefficients across channels. Each panel displays results for different low-frequency bands: Theta (left panels; shades of green), Alpha (middle panels; shades of red), and Beta (right panels; shades of blue). Results are shown for each SD condition, with values averaged across all samples per channel. **e** Root Mean Square Error (RMSE) was calculated to measure the variability in power dynamics between correct and wrong choice trials across SDs. RMSE values are shown in swarm plots, where each dot represents the deviation of a single channel averaged across all subsamples. **f** Average RMSE across channels for direct comparison of frequency bands, with coloured error bars indicating mean ± 1 SEM from panel e for both monkeys. The plot highlights a lower difference in power dynamics between correct and wrong trials for high-Gamma.

**Fig. 6.**
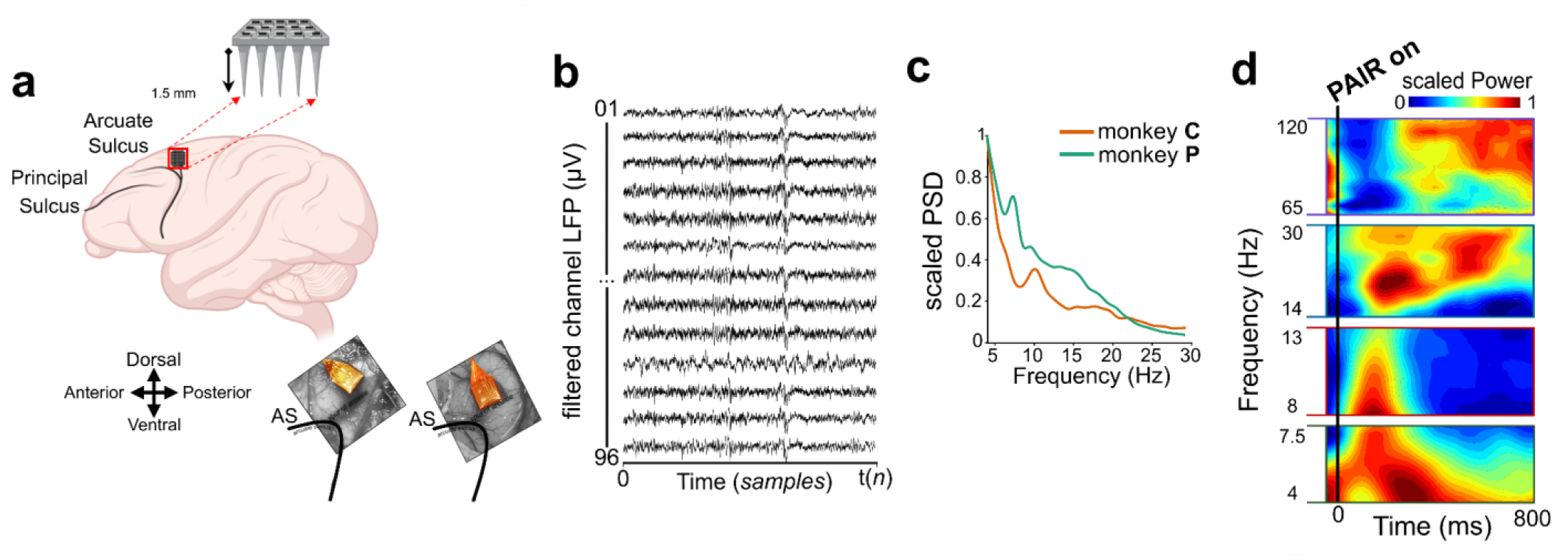
Mapping oscillatory power dynamics in the PMd using Utah Arrays. **a.** A macaque brain model highlighting the Arcuate Sulcus and Principal Sulcus, with the 96-channel Utah Arrays shown as a grid of black dots in the region of interest. A red square, with arrows pointing outward, the brain model, indicates the length of the electrodes (1.5 mm). Intraoperative images of the arrays implanted in the left hemisphere of both monkeys (monkey C left picture, monkey P right picture). **b** Filtered LFPs (in microvolts, µV) over time for a subset of channels, showing neural activity after filtering. The data captures a short period within a single session. **c** Power Spectral Density (scaled PSD) emphasizing variations in low-frequencies and spectral characteristics of two example channels for both monkeys. **d** Time-frequency representations of LFP data averaged across all channels and trials during a single session of monkey C (for SD=1). Vertical lines at the 0 ms mark PAIR onset, and a colour scale bar indicates decibels (dB) relative to the baseline period.

Power dynamics during wrong trials showed a disrupted pattern compared to correct trials, particularly in easier comparisons (SD4/5) (Fig. 5**b**). Unlike correct trials, where power scaled with comparison difficulty, wrong trials lacked consistency. To assess whether the different pattern of power dynamics in the wrong choice did not depend on the reduced number of trials, we conducted a control analysis aimed at balancing the number of trials in both correct and wrong trials. This analysis focused on low-frequency bands, as these oscillations serve as input samplers for the underlying computations. We first grouped correct trials from all sessions for each SD, ensuring an even distribution of trials on average (monkey C: *N*=865 ±179 trials; monkey P: *N*=851 ±70 trials). For each channel and SD (1-4/5), we randomly selected a number of correct trials equal to the number of wrong trials in the easiest condition (monkey C: *N*=23; monkey P: *N*=17 trials). The stability of difficulty level encoding was assessed using correlation matrices across 100 iterations per channel, SD, and frequency band during the modulated time periods (Fig. 5**c**). This allowed us to evaluate consistency in power dynamics, even with few trials, by averaging correlation coefficients across the different random samplings of trials in each lower frequency band. One-way repeated measure ANOVAs confirmed increased power stability in easier comparisons for both monkeys (Fig. 5**d**), with SD4/5 exhibiting significantly higher consistency (p < .001). Post hoc tests confirmed significant effects between SD4/5 and other SDs (p < .01). Once we established that the dynamics of oscillation were consistently modulated by rank difficulty level across different subsamples of correct trials and electrodes, we quantified, by the Root Mean Square Error (RMSE), the deviation in power during wrong trials from the identified modulation pattern in correct trials. The RMSE was computed for each channel, SD, and frequency band across all samples, ensuring an equal number of trials for comparison. Averaged RMSE values across samples for each channel were then compared across SDs (Fig. 5 **e-f**). This analysis revealed: **i**) a general increase in deviation of power dynamics across all SDs (all > 0); and **ii**) significantly greater deviations during the easiest decisions (SD4/5), compared to more difficult ones (SD1-3) (one-way repeated measure ANOVAs, both monkeys, all p < .001; see Supplementary Table S3). Post hoc tests confirmed significant differences between SD4/5 and other SDs (p < .01), further showing that power dynamics became more unstable during wrong trials in easier conditions. Interestingly, we observed that high-Gamma values, while preserving the overall pattern of increased deviation during easier decisions, were on average reduced in magnitude compared to those in lower frequencies bands (Fig. 5**f**). This may indicate that high-Gamma is less sensitive to the difficulty of rank encoding comparison than lower frequency bands, aligning with the hypothesis that item rank computation involves more the input stages rather than the output. Overall, this result suggests that the disruption in the consistency of power dynamics may fail to properly integrate mnemonic information, leading to both longer RTs and wrong choices.

## DISCUSSION

Two macaques were trained to solve a transitive inference task and construct mental representations of hierarchically organized perceptual stimuli that changed in each experimental session. Deciding on the task requires accessing this representation for matching each visual item perceptually available with its hierarchical representation. By studying neural computations subtending this task through the dynamic interplay of lower and higher LFP modulations, we provide evidence that PMd is a valid candidate for integrating perceptual and mnemonic information to guide decision-making.

We first show that the degree of difficulty in decision making was significantly influenced by the SD effect^13,15,24^, widely documented to depend on the access to a mnemonic schema where rank code proximity between items influenced monkeys’ capability to select the item assigned to reward delivery (Fig. 1**e-f**). Then we observed that LFP oscillations modulation reflected the degree of rank code comparison difficulty: lower frequency power decreased, negatively related to Accuracy (Fig. 2**a-c**, **e-g**), while high-Gamma power increased, positively related to Accuracy (Fig. 2**d, h**). These findings align with a documented dynamic interplay between low and high-frequency oscillations, which is thought to support the ongoing decision by the available information being integrated for achieving a final deliberation^16,25^. Here, it is plausible that the deliberation of decision, marked by high-Gamma power increasing, starts when the power of lower frequencies is sufficiently low. The present results display that this interplay depends on the difficulty of rank comparison. We further investigated whether the strength of the interplay between lower frequencies and high-Gamma differed. The result of this analysis detected different degrees of anticorrelation (Fig. 3**b**). The values of anticorrelation were particularly prominent between Beta and high-Gamma bands (Fig. 3**c**) and this anticorrelation was significantly longer in time (Fig. 3**d**). This suggests a different role, at least between the Beta band and the other two lower frequencies, in supporting the decision. Moreover, we observed that the number of channels recruited in these computations varied with the rank code comparison difficulty. During higher difficulty, there was an increase in lower frequencies power across a greater number of channels, whereas high-Gamma power was concentrated into fewer channels. As difficulty decreased, the reverse pattern emerged, suggesting that the transition from a more effortful cognitive state to a less demanding one is accompanied by a shift in the distribution of oscillatory power (Fig. 4**a-d**). These findings suggest the idea that the PMd adjusts its resources in response to the task’s demands, engaging more channels for complex computations and fewer for simpler tasks. Interestingly, we observed deviations from this pattern during wrong trials, particularly when the decision was easier (SD4/5) (Fig. 5**a-b, e-f**). This suggests that in easier trials, which are those requiring less integration effort, failures can still occur when the LFP signal consistently deviates from the appropriate pattern. Finally, a comparison of the deviation index of high-Gamma power between correct and wrong trials reveals a consistently lower deviation than that observed in lower frequency bands. This further supports the hypothesis that high-Gamma and lower frequencies bands play different functional roles and suggest that lower frequencies are more sensitive in processing properly incoming ranking-related information. Thus, our data suggest that the PMd, through its oscillatory activity and the interplay between lower frequency bands and high-Gamma, plays a critical role in decision-making based on the integration of perceptual and the paired abstract property stored in memory schemas. Specifically, the decrease in power of lower frequency bands appears to support the processing of memory-related information, while decision deliberation is associated with the increase of power of high-frequency oscillations.

The neuronal correlates of decision-making have been extensively studied using perceptual decision-making tasks, often based on the random dot motion paradigm. This task requires participants to discern the predominant motion direction of moving dots. Other studies have examined decision-making through static visual discrimination, such as orientation, contrast, or colour differentiation^2,3^. The independent variables in these tasks, such as dot movement coherence, line orientation similarity, and colour contrast, systematically vary task difficulty, allowing researchers to explore how sensory inputs are transformed into categorical judgments and the corresponding neuronal computations^2,26,27^. Similar principles have been extended to auditory^28^ and tactile stimuli^29^, advancing our understanding of perceptual decision-making mechanisms. A widely accepted model explains decision commitment as the temporal accumulation of evidence favouring one alternative, reflected in a ramping increase in spiking activity, with slope variations depending on perceptual decoding difficulty^1,30^. In contrast, the present study investigates decision-making based on implicit information embedded in perceptual stimuli, requiring deliberation beyond direct feature comparison. Here, difficulty arises from the implicitly ordered representation of item ranks and their relative values^13^, mentally represented within a geometric mental workspace^18^. Despite this distinction, spiking activity in the prefrontal and premotor cortex exhibits similar modulation patterns to those observed in perceptual decision-making^10,31^ and provides evidence of a possible common computational substrate.

Our study replicates these findings by showing that task difficulty modulates high-Gamma power, which reflects modulation in spiking activity^20,23,32^ and the neuronal computation output state. Additionally, low-frequency bands were also influenced, suggesting active information processing within the decision network. Given that this task cannot be resolved through simple perceptual analysis and considering the LFPs dynamics, we propose that PMd integrates and likely multiplexes memory-related information with perceptual input, potentially facilitating the ramping to threshold dynamics of high-Gamma bands.

Beta-band activity (∼13–30 Hz) in sensorimotor regions has traditionally been linked to motor preparation, decreasing before movement and increasing post-movement ^33^. However, Beta activity in the macaque^34^ and human^35^ premotor cortex also encodes categorical decision signals independent of motor response mapping. Our results provide further evidence of the role of Beta power in the PMd as a modulator in the transition between decision deliberation and motor action. Beta is implicated in the temporary processing of task-relevant information, facilitating endogenous retrieval of cortical representations^7–38^. These modulations are transient (several hundred milliseconds), emerging late in delayed tasks when information is refreshed for comparison rather than being maintained persistently^38^. Our results suggest that temporary information manipulation disengages once a comparison result is obtained, with the timing of disengagement varying based on the rank code comparison difficulty. Recent studies indicate that Beta power, linked to Beta burst changes, remains elevated for the duration of task-relevant information and decreases at the end of processing, often coinciding with increased spiking activity and high-Gamma power^16,25,39^. We observed an initial Beta power increase following PAIR onset, followed by a decrease modulated by rank code comparison difficulty, accompanied by an increase in high-Gamma power. More difficult comparisons required greater stimulus processing effort, leading to higher Beta power, likely reflecting increased cognitive demand in working memory-related brain regions^40^. Similar to the anticorrelation between Beta and high-Gamma, Theta (∼4–7.5 Hz) and Alpha (∼8–13 Hz) oscillations exhibit a complementary pattern in processing information flow. Theta supports memory processing, playing a critical role in encoding, retrieval, retention, and top-down control. In parietal, prefrontal, and premotor areas, increased stimulus load correlates with decreased Beta, Theta, and Alpha power, while high-Gamma power increases in response to reduced working memory load^40^. This modulation pattern suggests a functional trade-off between low and high-frequency oscillations in managing cognitive demands. Theta power is hypothesized to facilitate memory integration by coordinating information flow between the PFC and subcortical structures, such as the hippocampus^41^, which has been implicated in transitive inference tasks ^42,43^. A similar functional role may apply to the PMd, where Theta power has been shown to increase during memory encoding and task interruptions and decrease during retrieval^44^. This suggests that Theta oscillations could reflect the flow of rank-order representations from the hippocampus to cortical areas, with modulation corresponding to the difficulty of associating items with their relative ranks. Additionally, the inhibitory functional role of Alpha oscillations, acting through a “gate-keeper” mechanism, could support the decision-making process, where a decrease in Alpha power reflects the release of inhibition, enabling enhanced cognitive processing^45–47^. During working memory tasks, Alpha oscillations are thought to protect memory maintenance from distracting information^48,49^. In the present case, the greater overlap in item rank representations for smaller SDs likely led to a higher degree of interference, where the lower-ranking item exerted a stronger influence on the higher-ranking one. This interference effect could be reflected in the modulation of the observed Alpha power. Overall, the distinct functional roles of different frequency bands are in line with a model in which incoming visual and mnemonic information are integrated within a framework that gradually accumulates evidence favouring decision deliberation. In the present task, while low frequencies multiplex different pieces of information, the increasing activity of high-Gamma facilitates the maturation of decision deliberation. This perspective positions the PMd as a key player in decision-making, not simply associating perceptual information with motor output but also processing more abstract information stored in memory structures. Our findings suggest that the relative rank of visual items may be temporarily encoded in the PMd for the duration required to engage in a motor decision. This interpretation aligns with recent research highlighting the PMd as the locus where the rank order of items and motor decision-making overlap^50^. Here, we provide evidence that this encoding may be reflected in the dynamics of local field potentials.

A dominant role in complex decision-making and cognitive flexibility has long been attributed to the PFC, a brain region that receives inputs from various sensory and limbic areas and is connected to motor-planning regions^12,51^. However, several studies have demonstrated that both posterior, parietal, and premotor areas are also competent in managing variables that guide decision-making. Sensory and motor regions reflect decision formation processes that extend beyond encoding the sensory input or generating the motor output^4^. The distributed presence of neural activity encoding the decision variable may converge on the hypothesis that many brain structures form modular interconnected networks supporting different forms of decision-making^4,52^. The different areas compute several attributes characterizing the available options, converting them into a format suitable for threshold-based decision selection, which in motor-related areas emerge as motor plans^53,54^. Within this framework, motor-planning areas do not merely receive abstract decision variables from an amodal central module capable of decision making, as could be the PFC. Rather, they could be part of the network that flexibly transforms sensory signals into action plans^4^. In this distributed modular network, there is no clear parcellation of brain regions into those responsible for flexible control and others. However, there are gradients across brain structures, where some areas have more capacity to maintain task rules, flexibly adjust computations, and exert influence over other regions. Therefore, higher-order sensory and motor-planning cortical regions and subcortical areas that have access to both sensory and motor information could play significant roles in flexible behaviour, as earlier sensory areas could do with lower weight. The PFC lies at the top of this distributed network as the most flexible circuitry, but does not play a unique role as a control area^4,53,54^. This picture fits the comparable functional properties of prefrontal and premotor cortex^12,51,55^, as well as not strictly motor-related computations detected in premotor cortex^3,11,56,57^. The present work provides evidence on the PMd’s role in decision-making with a form of computation that combines perceptual (explicit) and mnemonic (implicit) information. This computational property confers to this brain area a higher degree of flexibility in adapting to the environmental variability, predicting future events without direct experience, which represents a characteristic that goes beyond matching the perceptual stimuli and specific motor actions. Looking at these properties from an evolutionary point of view, as the development of the PFC is phylogenetically recent, while event prediction and flexibility in behaviour are common requirements for many organisms^59–62^, and it is plausible that this form of computation exists in structures less complex than PFC^62^.

## METHODS

### Animals

Two male rhesus macaque (Macaca mulatta), monkey C (10 kg) and monkey P (9 kg), were trained for the study. The monkeys were co-housed in enriched cages and received a daily diet consisting of standard primate chow supplemented with nuts and fresh fruits. To ensure hydration, part of their daily water intake was provided as juice during experiments. Fluid restriction was used for motivation, with regular weight and health monitoring. All procedures complied with European (Directive 2010/63/EU) and Italian (D.L. 26/2014) regulations and were approved by the Italian Ministry of Health.

### Experimental design and Transitive Inference (TI) task

Experiments were conducted in a dark, acoustically insulated room. Items were displayed on a 17-inch LCD touchscreen monitor (800×600 resolution, 75 Hz refresh rate, 200 Hz sampling rate, Micro Touch, USA). Each session consisted of a learning phase and a test phase. In the learning phase (Fig. 1**a**), monkeys were trained to learn the rank hierarchy of six items (different among sessions) presented in adjacent pairs. They learned through trial and error to select the higher-ranked item, which was rewarded. During learning the monkeys acquired the rank relation between pairs of adjacent items in blocks of 100 trials. The monkeys repeated the block until the average accuracy achieved at least 60% of correct choices. In some sessions (*N*= 16) we used a chain-linking learning approach in which the monkeys learned the ordinal relation between the first half (A>B>C) and the second half (D>E>F) of the set and then liked the two portions by the pair comparison C>D (see^10^). Regardless of the learning procedure, the ultimate goal was to establish a clear understanding of the learned rank-ordered hierarchy among the six items (in the figure, color-coded and labelled ‘A’ to ‘F’ for simplicity). Each trial began with the appearance of a white dot (Central Target, CT) at the centre of the screen. Touching and holding the CT (within 5 s) triggered the pair presentation (PAIR on). After a variable delay (0.6–1 s, 200ms steps), CT disappeared, signalling the Go Signal (Go on), at which point monkeys could make their choice by reaching toward one of the two items (Fig. 1**b**). In the test phase, monkeys applied the learned hierarchy to solve inferential problems involving both adjacent and non-adjacent pairs (e.g., B>D). All 15 possible pairs were randomly presented, each repeated at least 18 times, with the higher-ranked item appearing equally on both sides (Fig. 1**c**). To prevent familiarity-based strategies, abstract black-and-white images (16°×16° visual angle) were used and changed daily. Item presentation and behavioural event collection were coordinated through Cortex’s freeware software package (https://nimh.nih.gov/), synchronized with neural recordings.

### Behavioural measures

We assessed whether monkeys’ ability to select the higher-ranked item depended on decision difficulty, quantified as the SD between items in their hierarchical mental representation. The 15 pairs were grouped by SD, with larger SDs indicating easier decisions and smaller SDs indicating higher difficulty (Fig. 1**c**). Behavioural performance was evaluated using Accuracy and Reaction times (RTs). Accuracy, defined as the probability of selecting the higher-ranked item, was calculated for each SD to assess performance across rank code comparison difficulty levels. RTs were measured as the time from the Go Signal to movement onset (Mov. on) toward the chosen item.

### Neural data collection and frequency band power

Following behavioural training, each monkey underwent surgical implantation of a 96-channel Utah Array (Blackrock Microsystems, USA) in the left PMd, using the Arcuate Sulcus as a landmark after the opening of the dura (Fig. 6**a-b**). Implantation was performed under sterile conditions with veterinary supervision. Post-surgical care included antibiotics and analgesics. Electrodes (1.5 mm length, 0.1–0.8 Ω impedance) were arranged in a 400-micron-spaced grid. Neural recordings began ≥1 week post-surgery.

Extracellular signals were amplified and digitized using a Tucker Davis Technologies system (Alachua, FL, USA) with a PZ2 preamplifier, capturing the unfiltered raw signal at a sampling rate of 24.4 kHz. Raw signals underwent baseline correction by subtracting the session-wide mean to remove offsets and slow drifts. This ensured that subsequent analyses focused on relevant oscillatory dynamics of interest. LFPs were extracted using a two-stage Butterworth filtering process: high-pass (1 Hz, 2nd order) and low-pass (120 Hz, 5th order). The resulting filtered LFP signal was down sampled to 1 kHz for 1 ms temporal resolution. To analyse oscillatory power dynamics, time-frequency decomposition was applied. Complex Morlet wavelet convolution was applied across 60 logarithmically spaced frequencies (4–120 Hz) with wavelet widths varying from 4 cycles (4 Hz) to 10 cycles (120 Hz), balancing the time-frequency resolution trade-off ^63^. Power spectral density (PSD) was estimated via the Welch method (1 s moving windows, 500 ms overlap on the entire raw channel signal) to define band-specific low-frequency ranges by identifying local peaks (two example channels from each monkey are depicted in Fig. 6**c**). Further refinement involved visually inspecting the spectral characteristics during the epoch of interest by examining time-frequency (TF) plots cantered around the PAIR on (Fig. 6**d**). Power was then computed relative to a pre-stimulus baseline (−500 ms to PAIR onset, ensuring at least 2 cycles of the lower frequency = 4Hz) and normalized using a decibel (dB) conversion formula.

### Inclusion criteria for sessions

Recording sessions were selected based on Accuracy being above 50% across all SDs, indicating performance above chance, to ensure data reliability. A total of 24 sessions met these criteria (monkey C, *N*=12 and monkey P, *N*=12), ensuring robust data for analyses.

## STATISTICS

### Behavioural analysis

To evaluate the relationship between the monkeys’ performance and decision difficulty (SD), we examined response accuracy and RTs. For each monkey, one-way repeated measures ANOVAs were conducted with SD as the factor. Results are summarized in Fig. 1**e-f**, with detailed session data in Supplementary Table **S1.**

### Correct choice power dynamics analysis

To assess whether oscillatory power during the test phase encoded item rank representation modulated by rank-code comparison difficulty (SD), we performed time-frequency analysis. We first averaged power across trials for each SD for the Theta, Alpha, Beta, and high-Gamma frequency bands. Then, we pooled the data from all sessions and combined all channels for each monkey. One-way ANOVA tests were performed at each 1-ms time point starting from PAIR onset up to 800 ms to identify significant power modulations, assessing power differences across SDs. Bonferroni corrections (α = 0.01) were applied to control for multiple comparisons. These SD-modulated periods were then used to calculate the average power across all channels for each SD and monkey. To confirm significant modulation of frequency power by rank-code comparison difficulty, one-way repeated measures ANOVAs were performed.

### Linear regressions

To evaluate the relationship between SD-modulated power changes and behavioural outcomes, linear regression was performed, using averaged SD-modulated power as the predictor variable and response accuracy as the predicted variable. The linear regression model was specified as: *Y* = β0 + β1*X* + ε where *Y* represents response accuracy, β*0* is the intercept, β1 represents the slope of the regression, X denotes the averaged SD-modulated power values, and ɛ is the error term.

### Cross-frequency correlations

To assess the interplay between each lower frequency and high-Gamma, cross-frequency Pearson correlation coefficients were computed between the power values at each time point in the delay epoch for each channel using data pooled across sessions and SDs (Sessions = 12 x SDs = 5, total of 60 power values). Power time series were generated for each frequency pairing (**θ–hγ**, **α-hγ**, and **β–hγ**). To evaluate differences in interplay magnitude among frequency pairings, one-way repeated measures ANOVAs were performed on maximum anticorrelation peaks values. To further evaluate the consistency of anticorrelation periods at the channel level, we quantified the duration of significant correlation periods and assess for differences across frequency pairing. LSD post hoc tests were performed to control for significant effects.

### Channel contribution

To assess individual channel contributions to SD-modulated power in the delay epoch (Fig. 2**a-d**), contribution ratios were calculated as the proportion of each channel’s power relative to the session’s total SD=1 power (used as a reference). This enabled session-by-session evaluations of channel recruitment across SDs. A threshold of 1.04% was established for significant contributions, derived from the assumption of equal power distribution across the array’s 96 channels, calculated as: *Contribution Threshold = (1/Number of Channels) × 100 = (1/96) × 100% ≈ 1.04%*.

### Wrong choice analysis

RTs for wrong choices were compared to correct trials by calculating a Difference Index (Wrong RTs - Correct RTs) for each SD. A two-way ANOVA (Monkey and SD) was performed to assess RT differences between conditions. A control analysis was performed by subsampling correct trials to match the number of wrong trials in the easiest decision conditions (SD4/5). Stability in correct trials power dynamics was evaluated using correlation matrices across 100 subsampling runs for each channel, SD, and frequency band. The subsamples by sum samples correlation matrices were calculated using the average power during the modulated time periods of the delay epoch for each subsampling run. One-way repeated measures ANOVAs were performed to assess differences in power stability of channels between SDs. To measure deviations in power dynamics between correct and wrong trials, for each frequency band, we computed the RMSE for each channel, SD, and frequency band. One-way repeated measures ANOVAs were used to compare RMSE values of channels across SDs to assess the consistency of power dynamics. LSD post hoc tests were performed to control for significant effects. All statistical analyses and data processing were conducted using MATLAB (www.mathworks.com, Release: 2022a), utilizing both built-in functions and custom-made scripts.

## Author contributions

E.B., and S.F.: Conceptualization, Supervision, Funding acquisition; S.F.: Resources; E.B. and I.B.M.: Investigation, Methodology, Data Interpretation; I.B.M.: Data curation, Formal analysis; I.B.M. and V.G.: Visualization; I.B.M., E.B., S.F.: Writing of original draft; V.G., S.R., G.B.: Writing – review & editing

## Acknowledgments

We are grateful to Dr. Valentina Mione for her significant contribution to the implementation of the experiment and the collection of data, and to Dr. Marta Andujar for her support and assistance with the pre-processing of the neural activity.

## SUPPLEMENTARY MATERIAL

**Fig. S1.**
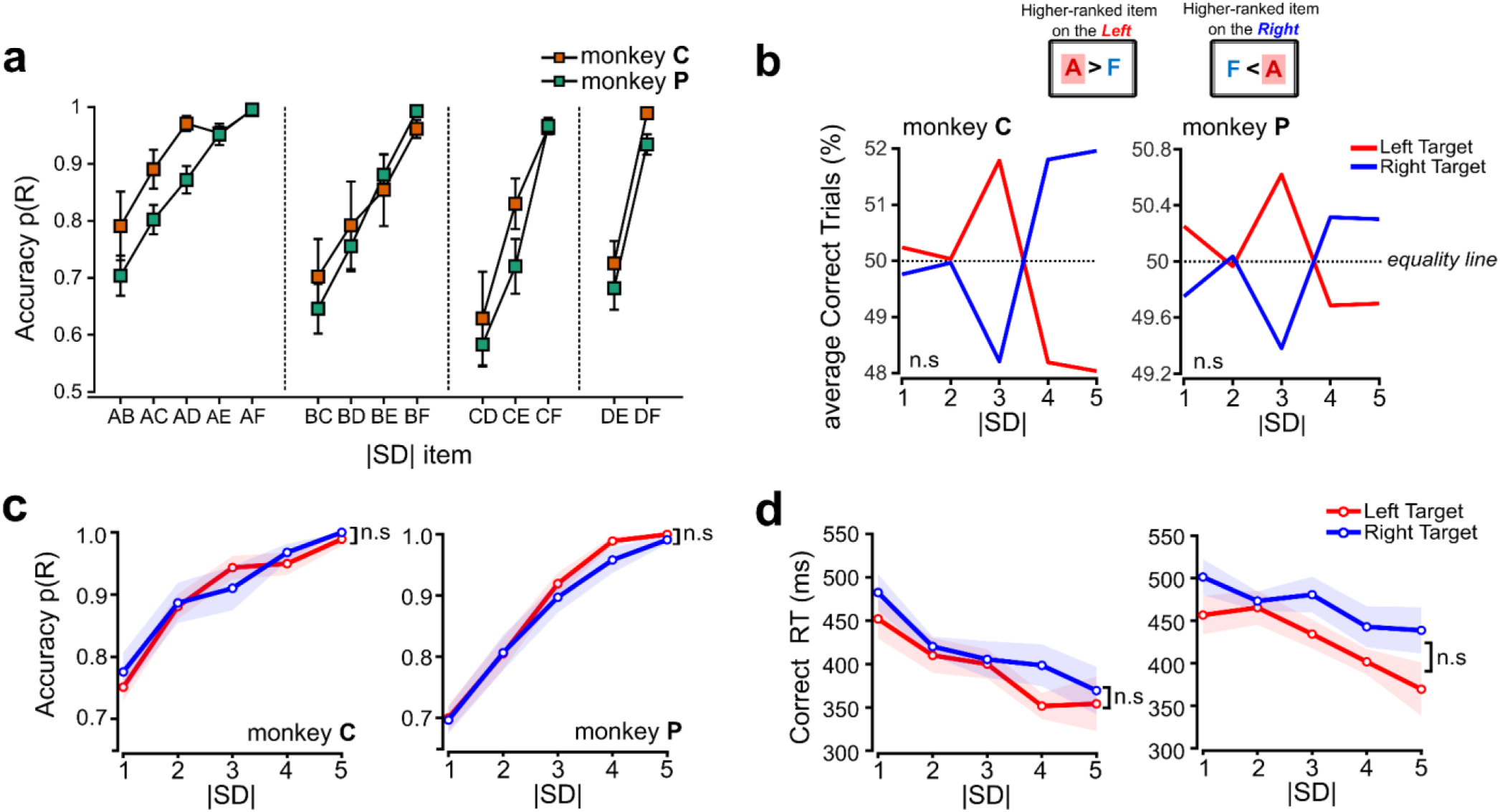
Performance in the TI task based on the spatial location of the higher-rank item and test of SD effect for individual items. Related to Fig 1**. a** Accuracy comparing each item separately with the other for each SD. Significant effect of SD emerged in all comparisons among items (monkey C: item A= slope + .047, p < .001; item B= slope + .084, p = .003; item C= slope + .167, p < .001; item D= slope + .263, p < .001; monkey P: item A= slope + .073, p < .001; item B= slope + .116, p < .001; item C= slope + .191, p < .001; item D= slope + .252, p < .001; linear regressions). Black error bars represent ±1 SEM. Despite item A being always the winning item, Accuracy did not reach 100% when presented, suggesting that rank code comparison difficulty (SDs) was likely driven by the overlapping representations in the hierarchical mental representation (see Fig. 1c). **b** To test for potential bias due to the spatial position of the target, we also examined whether monkeys showed a preference for one spatial location over the other in correct trials. Across sessions we calculated the percentage of correct responses made to either the Left (red line) or Right (blue line) side for each SD, equality line shows equal percentage 50%, with no significant bias across both spatial location (monkey C: F (1,11) = 0.23, p > .05; monkey P: F (1,11) =0.00, p > .05; two-way repeated measures ANOVA) and SDs (monkey C: F (4,44) =0.00, p > .05; monkey P: F (4,44) =0.00, p > .05). **c-d** Accuracy and Correct RTs were analysed to assess whether the spatial location of the higher-ranked item influenced performance. However, the spatial location did not significantly affect either Accuracy (monkey C: F (1,11) = 0.09, p > .05 and monkey P: F (1,11) = 0.73, p > .05; two-way repeated measure ANOVA) nor Correct RTs (monkey C: F (1,11) = 2.05, p > .05 and monkey P: F (1,11) = 2.13, p > .05; two-way repeated measure ANOVA), suggesting that spatial position within this representation is not a critical factor in monkeys’ performance.

**Fig. S2.**
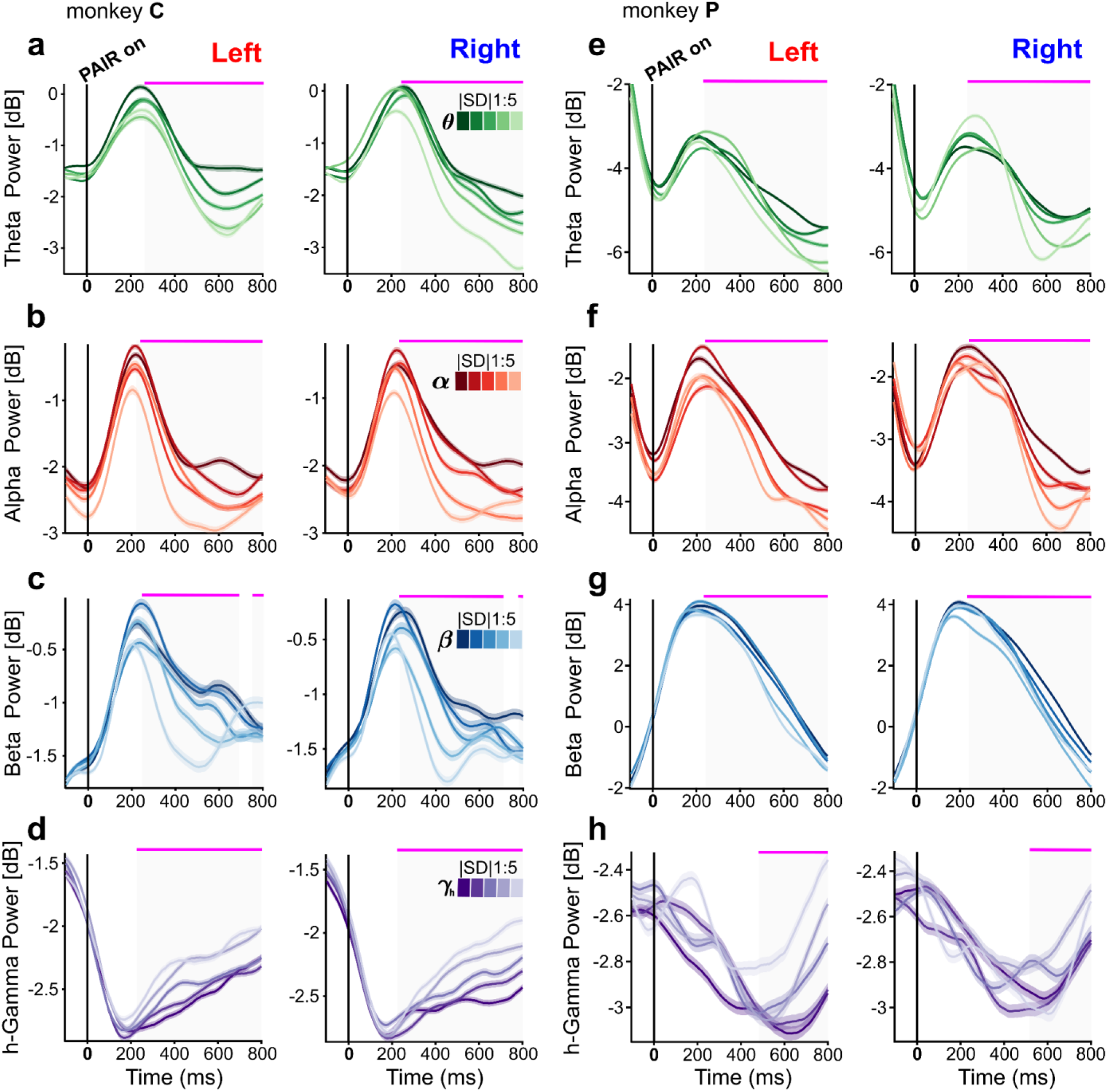
SD-related power changes aligned to PAIR onset separated by spatial position (Left or Right target) of higher-ranked item. Related to Fig 2**. a–d** monkey C and **e–g** monkey P. Left panels show trials where the higher-ranked item appeared on the left side; right panels show trials where it appeared on the right. Theta power (4-7.5 Hz; **θ** - shades of green), averaged over channels, *N*=1152 each, with shaded areas indicating ± 1 SEM. Significant differences in time are marked with a solid pink line and a grey background in each plot (one-way ANOVAs, p < .001, Bonferroni corrected with **α**= 0.01). The x-axis spans from −100 ms before to +800 ms after the PAIR onset. The y-axis indicates power magnitude in decibels (dB), normalized to the baseline period.

---

**Table S1.** Detailed behavioural outputs (Accuracy and Correct RTs) by session for each monkey. Related to. **Fig. 1f-g**

| sess<br>ion | monkey C |  |  |  |  |  |  |  |  |  | monkey P |  |  |  |  |  |  |  |  |  |
| --- | --- | --- | --- | --- | --- | --- | --- | --- | --- | --- | --- | --- | --- | --- | --- | --- | --- | --- | --- | --- |
|  | Accuracy |  |  |  |  | Reaction time |  |  |  |  | Accuracy |  |  |  |  | Reaction time |  |  |  |  |
|  | SD<br>1 | SD<br>2 | SD<br>3 | SD<br>4 | SD<br>5 | SD<br>1 | SD<br>2 | SD<br>3 | SD<br>4 | SD<br>5 | SD<br>1 | SD<br>2 | SD<br>3 | SD<br>4 | SD<br>5 | SD<br>1 | SD<br>2 | SD<br>3 | SD<br>4 | SD<br>5 |
| 1 | .79 | .93 | 1 | .88 | .93 | 518 | 511 | 423 | 428 | 428 | .68 | .84 | .86 | .97 | 1 | 597 | 582 | 587 | 567 | 541 |
| 2 | .84 | .85 | .98 | .96 | 1 | 512 | 450 | 427 | 404 | 397 | .74 | .76 | .98 | .97 | 1 | 461 | 462 | 462 | 441 | 415 |
| 3 | .73 | .92 | .98 | .93 | 1 | 490 | 436 | 429 | 465 | 610 | .72 | .93 | .91 | 1 | 1 | 387 | 473 | 404 | 377 | 344 |
| 4 | .75 | .93 | .96 | 1 | 1 | 371 | 342 | 324 | 277 | 249 | .75 | .77 | .96 | 1 | 1 | 478 | 436 | 437 | 400 | 352 |
| 5 | .67 | .71 | .77 | .90 | 1 | 441 | 429 | 386 | 405 | 388 | .70 | .80 | .93 | 1 | 1 | 530 | 487 | 439 | 432 | 369 |
| 6 | .68 | .71 | .71 | .90 | 1 | 448 | 412 | 424 | 411 | 364 | .77 | .95 | 1 | 1 | 1 | 616 | 613 | 530 | 458 | 418 |
| 7 | .77 | .92 | .89 | 1 | 1 | 399 | 380 | 372 | 317 | 259 | .68 | .79 | .91 | .97 | 1 | 527 | 523 | 466 | 412 | 474 |
| 8 | .70 | .92 | .95 | .95 | 1 | 510 | 447 | 457 | 435 | 357 | .65 | .71 | .84 | .86 | 1 | 452 | 428 | 452 | 455 | 497 |
| 9 | .62 | .90 | .95 | .98 | 1 | 492 | 412 | 495 | 351 | 645 | .63 | .73 | .84 | .96 | 1 | 425 | 419 | 410 | 380 | 306 |
| 10 | .82 | .95 | 1 | 1 | 1 | 461 | 374 | 316 | 311 | 315 | .74 | .71 | .94 | 1 | .98 | 462 | 368 | 408 | 370 | 383 |
| 11 | .88 | .88 | .94 | .98 | 1 | 508 | 390 | 389 | 388 | 303 | .62 | .78 | .78 | .93 | .95 | 404 | 420 | 453 | 394 | 395 |
| 12 | .84 | .93 | .95 | .97 | 1 | 436 | 390 | 394 | 310 | 315 | .64 | .83 | .90 | .98 | 1 | 403 | 410 | 423 | 372 | 347 |

**Table S2:** One-way repeated measure ANOVA results. Related to. **Fig 2 a-d**

|  |  | Frequency Bands | F-values<br>(4, 5755) | p-value |  |  | Frequency Bands | F-values<br>(4, 5755) | p-value |
| --- | --- | --- | --- | --- | --- | --- | --- | --- | --- |
| monkey C |  | Theta | 412.66 | **p < .001** | monkey P |  | Theta | 201.50 | **p < .001** |
|  |  | Alpha | 551.35 | **p < .001** |  |  | Alpha | 283.84 | **p < .001** |
|  |  | Beta | 581.19 | **p < .001** |  |  | Beta | 436.65 | **p < .001** |
|  |  | high-Gamma | 529.99 | **p < .001** |  |  | high-Gamma | 216.56 | **p < .001** |

**Table S3:** One-way repeated measure ANOVA results. Related to **Fig 5 e-f**.

|  |  | Frequency Bands | F-values<br>(4,380) | p-value |  |  | Frequency Bands | F-values<br>(4, 380) | p-value |
| --- | --- | --- | --- | --- | --- | --- | --- | --- | --- |
| monkey C |  | Theta | 111.79 | **p < .001** | monkey P |  | Theta | 163,79 | **p < .001** |
|  |  | Alpha | 114.06 | **p < .001** |  |  | Alpha | 282.82 | **p < .001** |
|  |  | Beta | 43.33 | **p < .001** |  |  | Beta | 283.52 | **p < .001** |
|  |  | high-Gamma | 72.23 | **p < .001** |  |  | high-Gamma | 174.37 | **p < .001** |

